# Matrix Softening Controls Stretch-Induced Cellular Memory and Fibroblast Activation

**DOI:** 10.1101/2022.10.12.511903

**Authors:** Yuan Hong, Xiangjun Peng, Haomin Yu, Mohammad Jafari, Delaram Shakiba, Yuxuan Huang, Chengqing Qu, Ermia E. Melika, Andrew K. Tawadros, Aliza Mujahid, Jacob Sandler, Kenneth M. Pryse, Justin M. Sacks, Elliot L. Elson, Guy M. Genin, Farid Alisafaei

## Abstract

Mechanical stretching of living tissues can activate long-lived changes in tissue cells such as fibroblasts, increasing their contractility and initiating phenotypic transformations. Increased mechanical stimulus typically leads to monotonically increasing activation of fibroblasts cultured in 2D, but activation levels are difficult to predict for cells in 3D fibrous tissues, leading to variable outcomes in procedures such as skin grafting. Here we report that the source of this variation is cell-extracellular matrix (ECM) interactions and their variation with the duration and magnitude of applied stretch, and present a model that can predict the degree to which stretch will either increase or decrease long-term activation levels of fibroblasts cultured within a stretched, three-dimensional collagen matrix. Combining experimental and mathematical approaches across multiple scales, we show that the viscoplasticity of the ECM regulates this nonmonotonic, long-term cell activation. Results demonstrate that feedback between cell and ECM determines how cells retain memory of mechanical stretch.

## Introduction

Soft tissues generally exist in a state of homeostatic tension.^1^ During development, wound healing, and surgical procedures such as skin grafting, wound closure, and flap reconstruction, the homeostatic set-point can shift, potentially leading to complications such as fibrosis, hypertrophic scars, keloid scars, and failed skin grafting. These outcomes are difficult to predict, leading to significant patient burden. For example, failed skin grafting is a devastating and unpredictable potential outcome for those of the 8.2 million chronic wound patients in the US each year whose chronic wounds are treated with split-thickness skin grafts.^2,3^ For these patients, chronic wounds arising from burns, diabetic ulcers, skin cancer surgery, or infection require surgical grafting to protect the wound from the environment and pathogens.^4–8^ The skin graft consists of epidermis and a thin variable layer of dermis from a donor site that is harvested (Fig. S1A (i)), meshed with an array of slits (Fig. S1A (ii)), stretched (Fig. S1A (iii)), and then transplanted over a larger wound site (Fig. S1A (iv)).^9–12^ Skin grafts placed on wound sites contract due to forces generated by dermal fibroblast cells from the donor site that are transferred within the graft. The graft can fail if this long-term contraction is either too great or too small, leading to complications ranging from poor aesthetic outcomes to major functional limitations such as scar contracture that impairs the function of joints such as the ankle, axilla, elbow, and wrist.^13– 16^ However, the level of this critical long-term contractility currently cannot be predicted. We hypothesized that levels of long-term graft contractility arise from activation or deactivation of graft fibroblasts that can be predicted from the magnitude and duration of the tensile strains that are transmitted to them through the extracellular matrix (ECM).

The rationale for this hypothesis is the apparent propensity of many cells to seemingly remember their previous mechanical conditions through changes to gene or protein expression that persist over physiologically relevant timescales.^17–23^ An example is mesenchymal stem cells cultured on two-dimensional (2D) substrata, wherein memory of substrate stiffness affects subsequent cell differentiation,^18–20^ and can persist even after cells are moved to a new substrate of different stiffness.^21,22^ Cyclic dynamic stretching, as occurs in development, also induces mechanical memory in stem cells, but static stretches do not.^24^ Fibroblasts are known to remember the stiffness of 2D substrates they encounter, with memory of relatively stiff substrates persisting in lung fibroblasts for weeks, even after cells are returned to softer substrates.^17^ In fibroblasts, this memory of a stiff environment manifests as elevated contractility and as activation, wherein fibroblasts express myofibroblastic gene and protein expression signatures that are hallmarks of wound healing and fibrotic disease.

This motivated us to predict that such mechanosignaling may have sustained effects in the form of long-term activation or deactivation of fibroblasts. Forces on fibroblasts are often in the form of long-term static stretching or compressing of the tissues in which they reside.^25,26^ In the case of skin grafting, these forces are tensile and are known to vary with the direction of skin stretching relative to the “Langer’s lines” that define cell polarity and are associated with surgical incision directions that reduce scarring.^27–30^ However, it remains unknown whether fibroblasts can develop and maintain mechanical memory under static loading. Furthermore, these external and long-term static forces are transmitted to fibroblasts through their three-dimensional (3D) extracellular environment whose physical properties are known to change with the magnitude and duration of the external loading.^31–33^ Therefore, we studied whether fibroblasts can develop and maintain mechanical memory under static loading within physiologically relevant 3D extracellular matrices (ECMs).

## Results

### Sustained stretch of tissues over a clinically relevant range of strains leads to dramatically different homeostatic states depending upon strain magnitude and duration

To quantify the effect of sustained stretch on fibroblast activation, we studied the contraction of synthetic tissues containing human dermal fibroblasts and type I collagen (Fig. S2). Thin (∼0.1 mm thickness), ring-shaped specimens formed as the fibroblasts contracted the collagen onto the central cylinder of an annular Teflon mold over two days (Fig. 1A). Thin specimens were chosen to represent certain conditions of split-thickness skin grafting, with reduced graft thickness often correlating with excessive contractility.^13^ The resulting ring-shaped tissues were placed on a uniaxial stretching apparatus (Fig. 1B) and subjected to a prescribed stretching regimen (Fig. 1C). Tissues were stretched with directions, durations, and magnitudes of stretch chosen to represent split-thickness skin grafts. The circumferential direction of loading coincided with the direction of cell alignment (Figs. 1B,D), representative of the stretching of skin grafts along Langer’s lines (Fig. S1C). Tissues that were not in the unstretched control group were stretched with strains of 5% (“low”) or 30% (“high”), representative of average strains associated with the range of typical skin grafts (Fig. S1E). All tissues were stretched for either one hour or 24 hours, representative of different environments where a skin graft can reside on the human body in a static or dynamic wound environment such as over an immobile (e.g., trunk) or mobile (e.g., joint) recipient bed.

**Figure 1.**
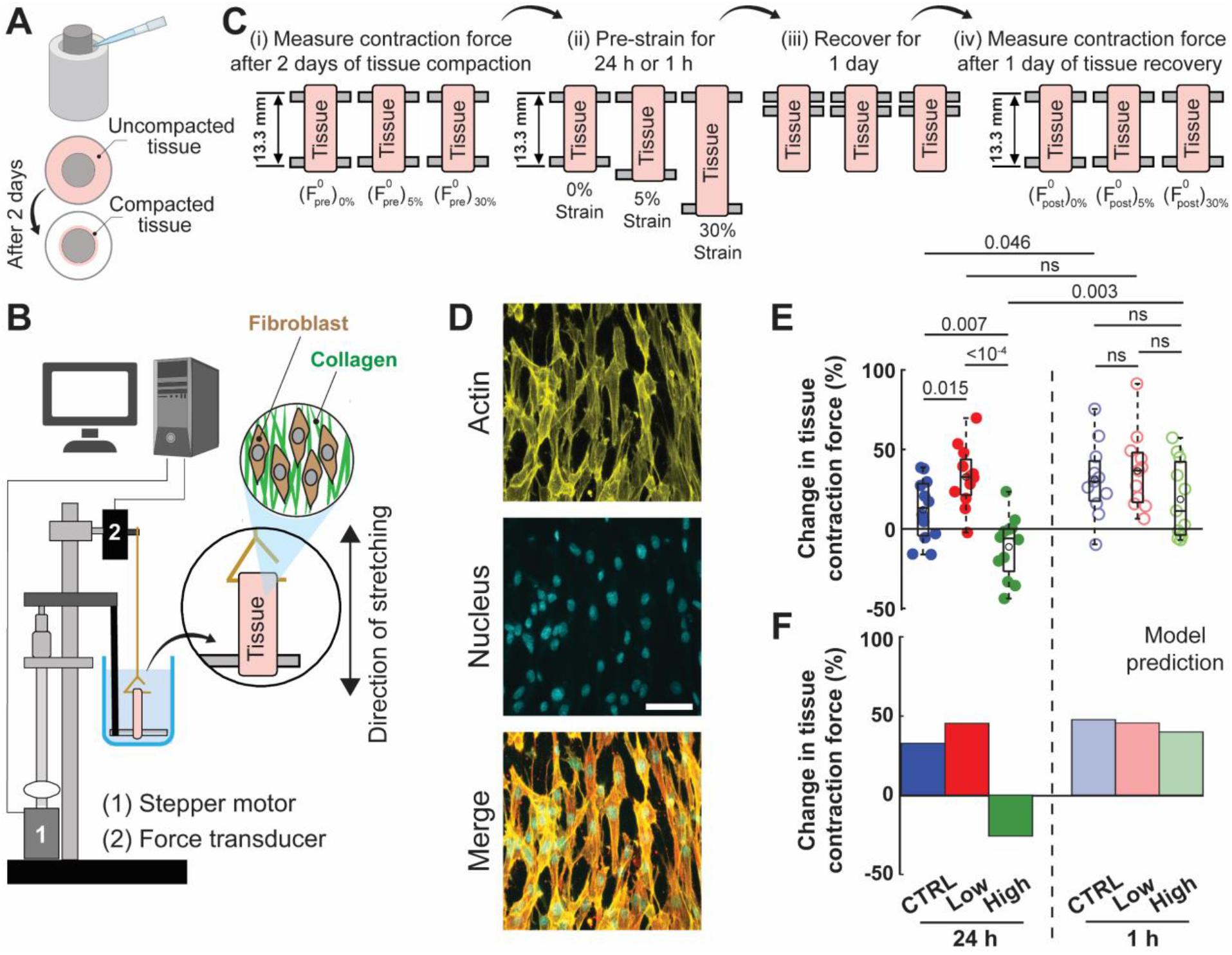
Experimental observations and computational simulations of the effects of sustained stretch on tissue contractility. (A) Ring-shaped tissue constructs consisting of human dermal fibroblasts and type I rat tail collagen were removed from molds after two days. (B) Tissues were placed on a force measurement device. (C) (i) The isometric contractile force (*F*^0^)_pre_ was measured in the reference configuration (inner perimeter of tissue equaling the inner circumference of the mold’s central cylinder). (ii) Tissues were strained 0% (CTRL), 5% (low), or 30% (high) and held for 24 h or 1 h, then (iii) allowed to relax for 1 day, (iv) after which post-stretch isometric force (*F*^0^)_post_ was measured in the unstretched reference configuration. (D) The cellular alignment within the tissues mirrored patterns observed along Langer’s lines *in vivo*. Tissues were stretched along the direction of cell alignment. Scale bar = 50 μm. (E) The change in the tissue contractile force was determined as Φ= ((*F*^0^)_post_ −(*F*^0^)_pre_)/(*F*^0^)_pre_ . Short-duration pre-straining had no effect on Φ, but a biphasic response was observed for long-duration pre-straining, with tissue contractility dropping following high strain (*n* = 12 and 11 for the long-duration and short-duration pre-straining, respectively.) (F) Our computational simulations predicted these results.

For each tissue, we first measured the isometric tissue contraction force (*F*^0^)_pre_ immediately after placing the tissue on the stretching apparatus on day 2 (Fig. 1C(i)), then “pre-strained” the tissue by 0% (control), 5% (low), or 30% (high) for either 24 hours (“long duration”) or 1 hour (“short duration”) (Fig. 1C(ii)). The tissues were then allowed to relax and return to the unstretched baseline length for one day (Fig. 1C(iii)), after which the isometric tissue contraction force (*F*^0^)_post_ (Fig. 1C(iv)) was measured in the unstretched reference configuration. The change in contraction force associated with the strain treatment, Δ*F*^0^ = (*F*^0^)_post_ −(*F*^0^)_pre_, was recorded.

For all three specimens stretched for 1 hour, Δ*F*^0^/(*F*^0^)_pre_ was positive, with the increase in contractility independent of stretch (Fig. 1E). However, tissues stretched for 24 hours showed a biphasic trend with respect to strain treatment: for tissues subjected to low stretch, Δ*F*^0^/(*F*^0^)_pre_was positive and maximum, while for tissues subjected to high stretch, the mean Δ*F*^0^/(*F*^0^)_pre_ was negative and significantly lower (*p* < 0.0001, Fig. 1E).

### The time- and strain-dependent properties of the ECM regulate changes to cell homeostasis and tissue contractility

To identify the sources of this unexpected, bimodal behavior, we separated the active and passive mechanical responses of the tissue specimens. After completing the 24 h of tissue recovery (Fig. 1C(iv)), viscoelastic stress relaxation testing was performed on each tissue, in which tissues were stretched by an amount ΔX=0.66 mm and held at that length while the isometric force was recorded (Fig. 2A(v)). The force rose to a peak during the stretching, and the subsequent isometric force then relaxed to an asymptotic value over time (Fig. 2B). The difference Δ*F* between the pre-test isometric force (force measured at 0 s) and the force measured after 600 s of relaxation was recorded, and the long-term stiffness Δ*F*/Δ*X* was calculated (Fig. S3). Tissues were subsequently returned to their reference lengths, treated with the actomyosin contractility inhibitor cytochalasin-D,^34^ then allowed to relax for one hour prior to repetition of the viscoelastic relaxation test to obtain the passive response of the tissues (Fig. 2A(vi-vii)). Cytochalasin-D treatment eliminated active contraction at the start of the relaxation test and reduced the peak force, but had little effect on the rate of long-term stress relaxation (Fig. 2B), as observed by others.^35^

**Figure 2.**
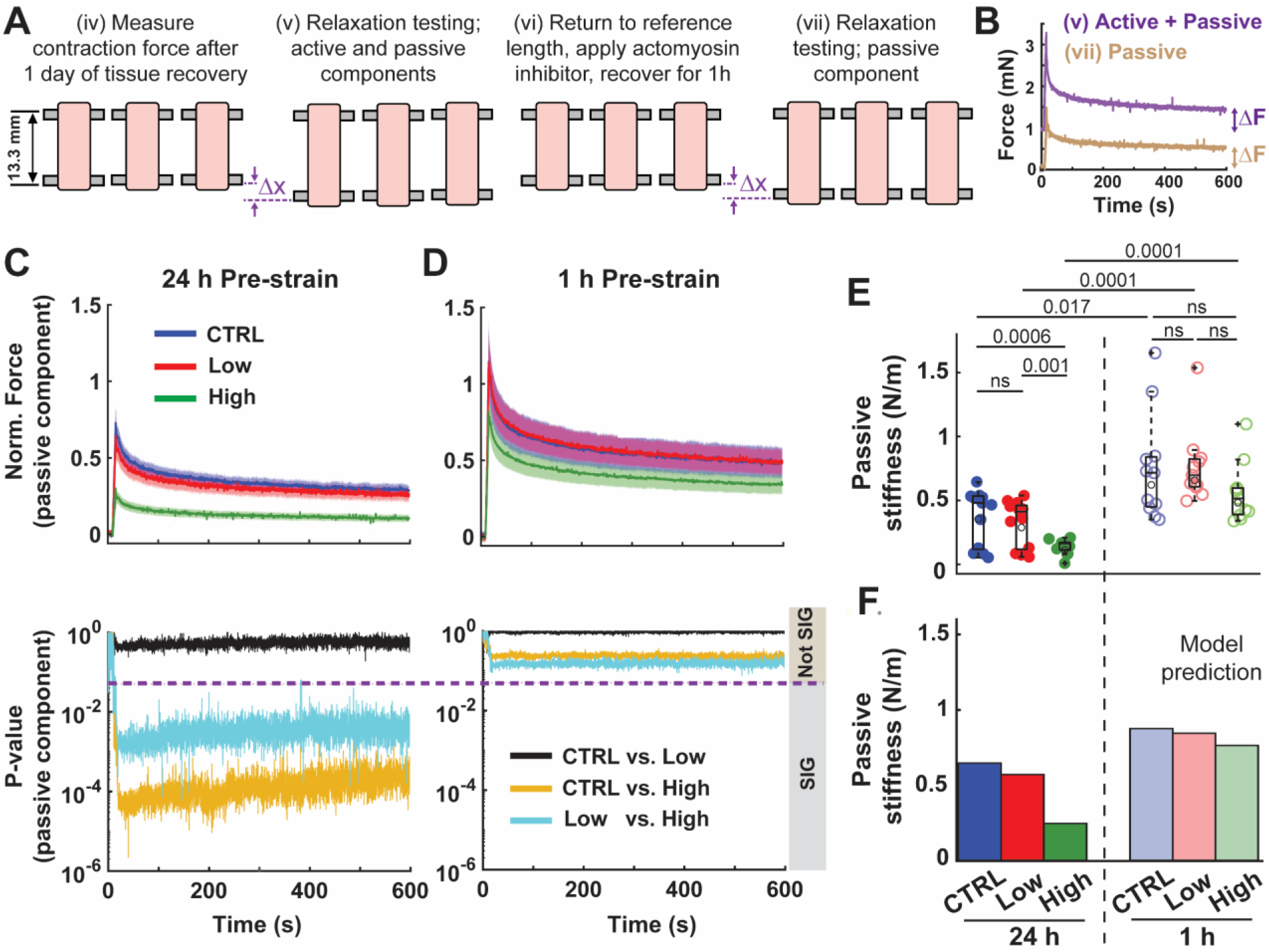
Experimental observations and computational simulations of the effects of sustained stretch on passive responses of tissues. (A) Effects of pre-strain magnitude and duration on the long-term passive responses of tissues were quantified via stress relaxation tests after the completion of tests described in Fig. 1. Isometric stress relaxation tests were conducted on the tissues (v) and then repeated (vii) with actomyosin contractility eliminated using cytochalasin D. (B) Representative stress relaxation responses of tissues before (purple line) and after (brown line) disruption of actomyosin contractility. Long-term stiffness was determined as Δ*F*/Δ*X*. (C-E) Disruption of actomyosin contractility revealed significant softening in tissues pre-strained for 24h, with softening further increasing with tissue pre-strain magnitude. On the contrary, pre-straining of tissues for 1 h did not cause permanent changes to long-term ECM stiffness (*n* = 13 and 11 for the 24h and 1h groups, respectively). In C and D, the solid lines and the shaded areas represent the mean and standard error, respectively. (F) Passive stiffnesses predicted by the active chemo-mechanical tissue model agreed with experimental observations.

Our results showed that tissues pre-strained for 24 h (long duration) had significantly softer ECMs compared to tissues pre-strained for 1 h. Furthermore, while the long-term ECM stiffness of tissues pre-strained for 1 h remained independent of pre-strain level, the long-term ECM stiffness of tissues pre-strained for 24 h varied with pre-strain level, with the ECM of tissues pre-strained at 30% for 24 h significantly softer than the ECM of tissues pre-strained at 0% or 5% (Figs. 2C-2E). Thus, pre-straining of tissues for 1 h did not cause permanent changes to long-term ECM stiffness, with the ECM showing full elastic recovery, but pre-straining of tissues for 24 h led to ECM softening, with the degree of softening affected by the pre-straining magnitude.

### Time- and strain-dependent ECM behavior regulates the stress levels that cells experience

To understand mechanisms underlying the effects of pre-strain duration and magnitude on tissue contractility (Fig. 1E-F), we linked the passive and active responses of the tissues through a computational model. This included an active chemo-mechanical tissue model (Fig. 3) coupled with a cell signaling network model (Fig. 4D). The tissue-level model predicted how forces reach cells through the ECM, and accounted for how ECM properties change with magnitude and duration of stretching (SI 1). The cell-level signaling model predicted how these forces impact long-term, time-dependent behavior of cells (SI 2).

**Figure 3.**
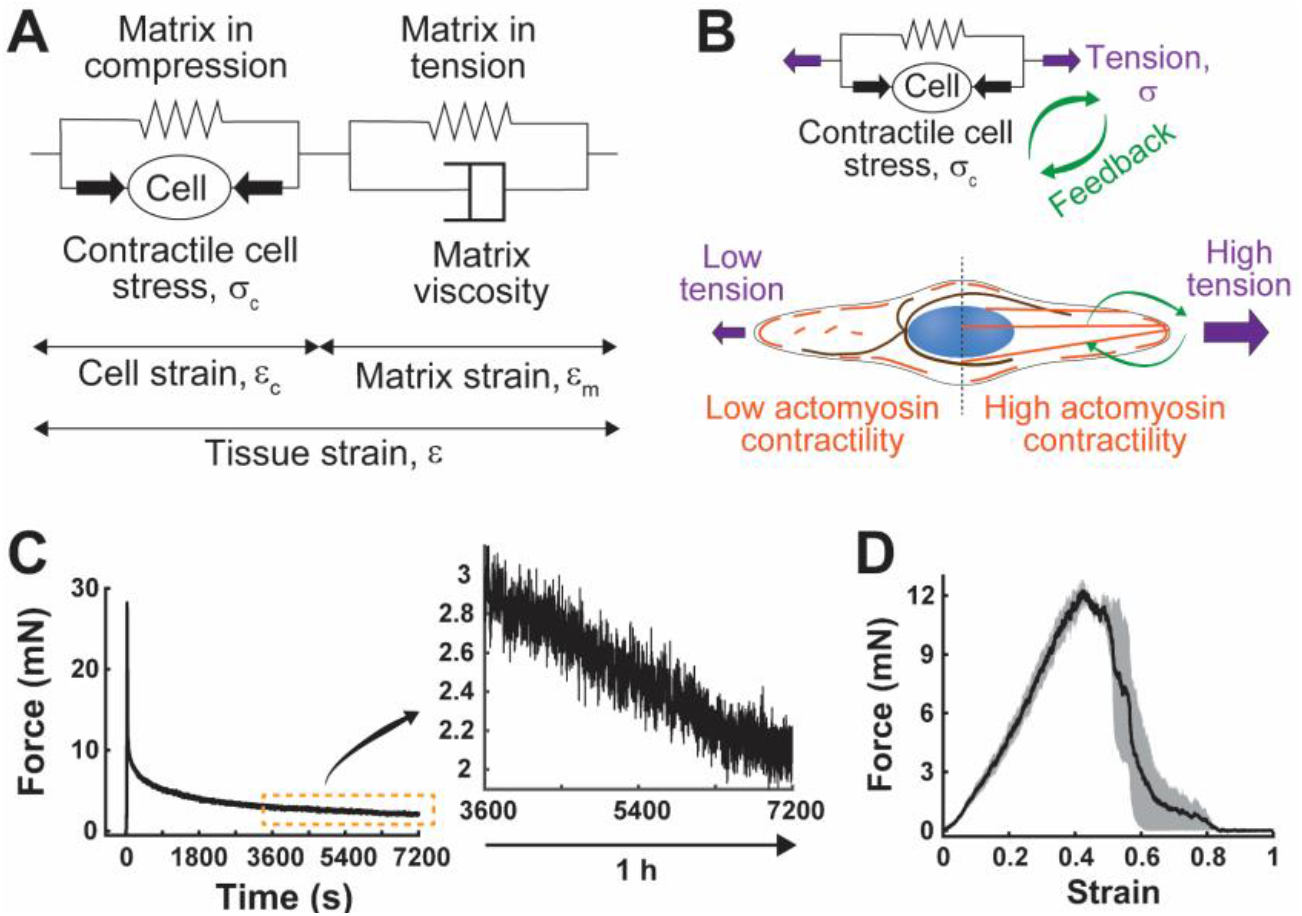
Time- and strain-dependent ECM mechanics. (A) A one-dimensional representation of the active chemo-mechanical tissue model composed of an active force-generating element (cell) connected to two passive elements. In the absence of external strain (ε = 0), one element represents regions of the ECM under compression due to cell contraction, and the other represents regions of the ECM in tension. The model represented the following three experimental observations. (B) First, actomyosin contractility and thus cell contractile force increase with tension. (C) Second, ECM exhibits long-term relaxation under tension. (D) Third, long-term ECM stiffness varies with ECM strain magnitude. The initial increase in the slope of a force-strain curve is followed by a softening region at larger strains (*n* = 3, the solid line and the shaded area represent the mean and standard error, respectively).

**Figure 4.**
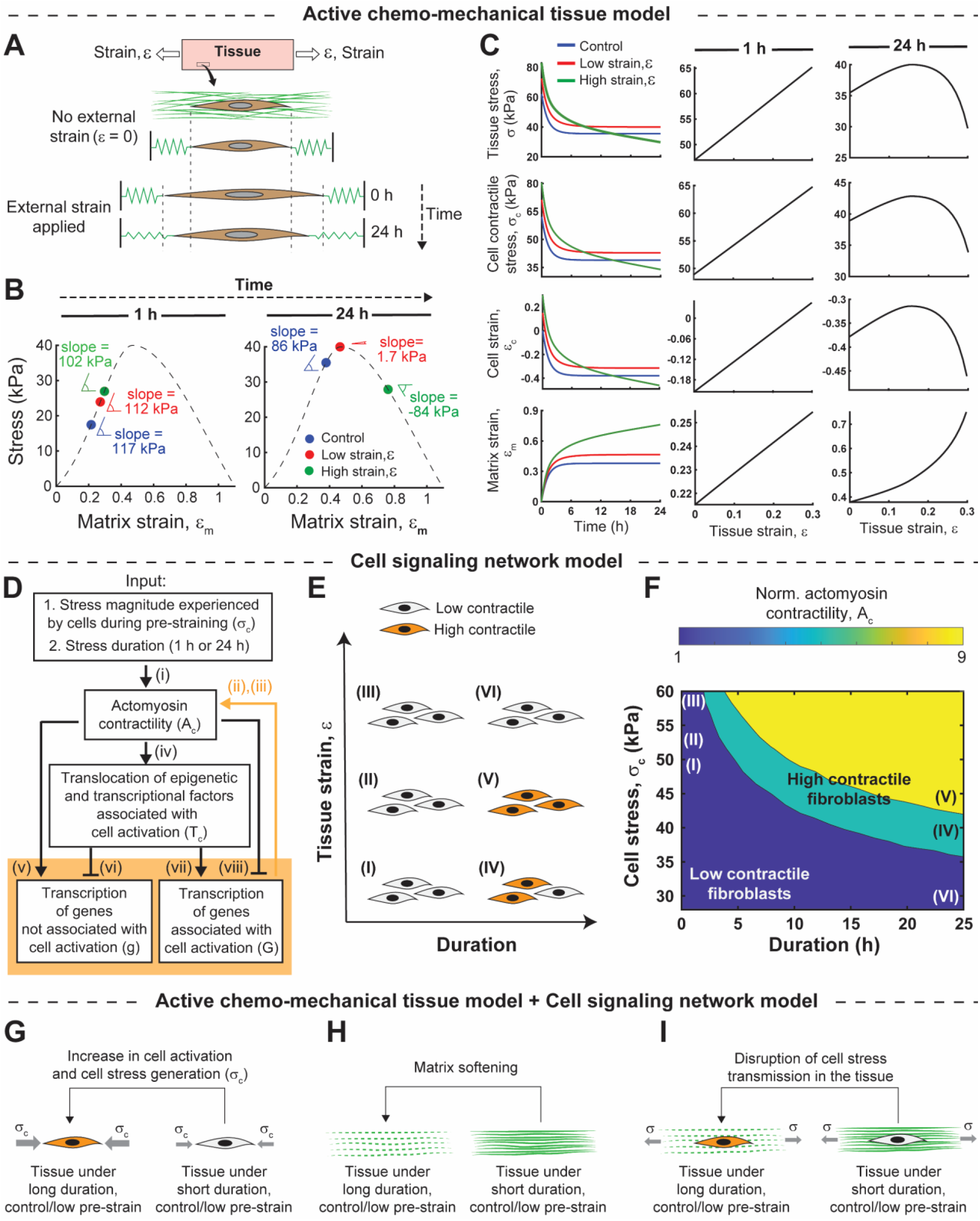
Stresses experienced by cells in a 3D ECM are regulated by time and strain-dependent ECM mechanics. (A) In stretched tissues, the strain ε_m_ in the ECM increases over time due to ECM viscous nature. (B) Over longer stretches (e.g., 24 h), ε_m_ can be stretched beyond the point at which the ECM softens. Dashed lines: average passive constitutive behavior of tissues from three experiments (cf. Fig. 3D). (C) For tissue strain magnitude, ε, applied over 24 h at a strain sufficiently high to cause this ECM softening, the model predicts that both the stress generated in the tissue, *σ*, and the cell stress, *σ*_*c*_, drop significantly. On the contrary, shorter durtations of stretch (1 h), are insufficient for creep to lead to ECM softening, so that both the stress generated in the tissue *σ* and the stress that cells experience *σ*_*c*_ are predicted to increase monotonically with the tissue strain magnitude ε. (D-F) Combining these cell-level predictions with a cell signaling model enabled prediction of a phase diagram that captured the biphasic activation of cells. (G-I) The integrated model thus can predict the seemingly contradictory observations in activation of cell contractility and tissue contraction force as a function of the magnitude and duration of tissue stretch.

The chemo-mechanical tissue model consisted of an ECM model and a cell model endowed with the ability to generate active forces (Fig. 3A). The ECM model was based upon two experimental observations. The first was that the ECM’s mechanical properties exhibit long-term, time-dependent changes under sustained tension. This hypothesis was supported by passive stress relaxation tests like those of Figs. 2C-D, but performed at a higher stretch (ΔX=4 mm) and over a much longer timescale (Fig. 3C). Results showed that the passive component of the tissue response continued to decrease with time, resulting in more than a 30% drop in matrix stress over 1 h (Fig. 3C). This ECM behavior was modeled by adding a viscous element to the ECM response in tension (Fig. 3A, SI 1.1).

The second was that ECM stiffness changes with magnitude of tensile ECM strain. To test this hypothesis, we stretched tissues treated with cytochalasin-D at a slow strain rate of 3.11 μm/s to minimize the ECM viscous effect. Experiments showed strain-stiffening up to a peak value, followed by a softening region (Fig. 3D). This was modeled by incorporating this experimentally measured stress-strain relationship into the constitutive model of the ECM element in tension (Fig. 3A, SI 1.1).

The cell model was based on the observations that actomyosin contractility and thus cell-generated contractile forces increase with cell tension, including tension arising from external tensile forces and from ECM resistance to cell contraction (Fig. 3B). This is supported by data from the literature, including our previous work, showing increased actin stress fiber polymerization in the direction of an applied stretch for cells in a 3D ECM,^36,37^ and data from 2D studies showing increased actomyosin contractility and cell force generation in response to higher matrix stiffness.^38–40^ This was modeled by incorporating cell-generated contractile stress, *σ*_*c*_, that increased with the tissue stress, *σ* (Figs. 3A,B and SI 1.3).

In a tissue stretched and held isometrically at strain ε, the ECM is stretched in tension by both cell-generated contractile stresses (*σ*_*c*_) and the external stresses applied to maintain the tissue strain of ε (Fig. 4A). However, due to the viscous nature of the ECM, the tensile strain in the matrix, ε_m_, increases with tissue stretching duration when the tissue is stretched isometrically (Fig. 4A). As a result, for long stretching durations (e.g., 24 h), the strain in the ECM can locally exceed the strain associated with peak stress in Fig. 3D. This effect could be seen in our simulations: after 1 h of isometric tissue stretching, the ECM strain stayed sufficiently low that the slope of the stress-strain curve (*E*_tangent_) was independent of tissue strain magnitude ε, but after 24 h, the ECM strain reached or passed the peak so that *E*_tangent_ dropped and varied with ε (Fig. 4B). To verify the model predictions against our experiments in Fig. 2E, we calculated the ECM stiffness (*k*), and found the model prediction to be in close agreement with those from our experiments (Fig. 2F, SI 1.6).

The model also provided a mechanism for how changes in ECM stiffness affect the tension generated in the tissues, *σ*, and subsequently the cell stress *σ*_*c*_ (Fig. 4C). For 1 h of tissue stretching, the ECM strain remained below the value associated with peak stress so that the stress generated in the tissue *σ* increased monotonically with the tissue strain magnitude ε (Fig. 4C). However, for 24 h of tissue stretching, the strain in the ECM passed the peak, softening the ECM. This effect increased with tissue strain magnitude ε such that for high ε, the stress in the tissue, *σ*, dropped, which in turn decreased cell-generated stress *σ*_*c*_ (Fig. 4C). Taken together, results showed that time- and strain-dependent changes in the ECM stiffness alter the stress generated in a stretched tissue, which in turn affects the stress level that cells experience.

### Increasing magnitude and duration of stress can increase or decrease the long-term activation level of cells

Inspired by a model of stem cell differentiation,^41^ we next developed a signaling network model (Fig. 4D, SI 2) and coupled it to the active chemo-mechanical tissue model to interpret data for how the magnitude and duration of the stress on cells in stretched tissues affected the long-term activation of cells that persisted after the tissues were unloaded. The signaling network model incorporated the following cascade of biological processes.

First, tensile stress promoted cell actomyosin contractility through pathways such as Rho-Rock or activation of mechanosensitive ion channels (Fig. 4D i).^42–45^ While excessive cell contraction can cause detachment of the cell from the extracellular matrix,^46^ thereby switching cells to a transcriptionally less active state (Fig. 4D v,viii),^25^ modest increases in cell contraction lead to translocations of transcriptional and epigenetic factors (e.g., MKL and YAP/TAZ) to the nucleus (Fig. 4D iv),^40^ which in turn promotes transcription of genes associated with actomyosin contractility and cell activation (Fig. 4D vi,vii).^25,47^ Finally, expression of these genes increases cell actomyosin contractility which is captured in our model through a feedback loop (Fig. 4D ii,iii).

The input to the signaling network model was the time variation of cell stress during tissue stretching, as predicted by the active chemo-mechanical tissue model (Fig. 4C). The output was the level of cellular actomyosin contractility that persisted after releasing the tissues. Note that cell stress *σ*_*c*_ was relatively high in tissues pre-strained for 24 h, displaying a monotonic increase with tissue strain ε (Fig. 4C). Conversely, cell stress was relatively low in tissues pre-strained for 1 h and exhibited a biphasic trend with tissue strain (Fig. 4C). The result was that cells in tissues pre-strained for 24 h with relatively low stretch were predicted to retain a higher state of activation than cells in tissues pre-strained for 24 h with relatively high stretch (Figs. 4E-F, 5C). Cells in tissues pre-strained for 1 h, even for cases of high tension magnitude, were not predicted to retain high levels of activation (Figs. 4E-F, 5C).

These cell-level model predictions were counterintuitive, given the diametrically opposite experimental observation at the tissue level, with pre-strained tissues for 24 h having significantly lower contractility than tissues pre-strained for 1 h. To verify the cell-level model predictions, we fixed tissues one day after tissue unloading and stained them for F-actin (Figs. 5A-B) and α-SMA (Fig. S4), which are well-known indicators of the level of fibroblast activation. Results were consistent with model predictions, and supported our hypothesis that long-term activation of cells that reside within a tissue is modulated by the way that force is conducted to these cells through the ECM.

**Figure 5.**
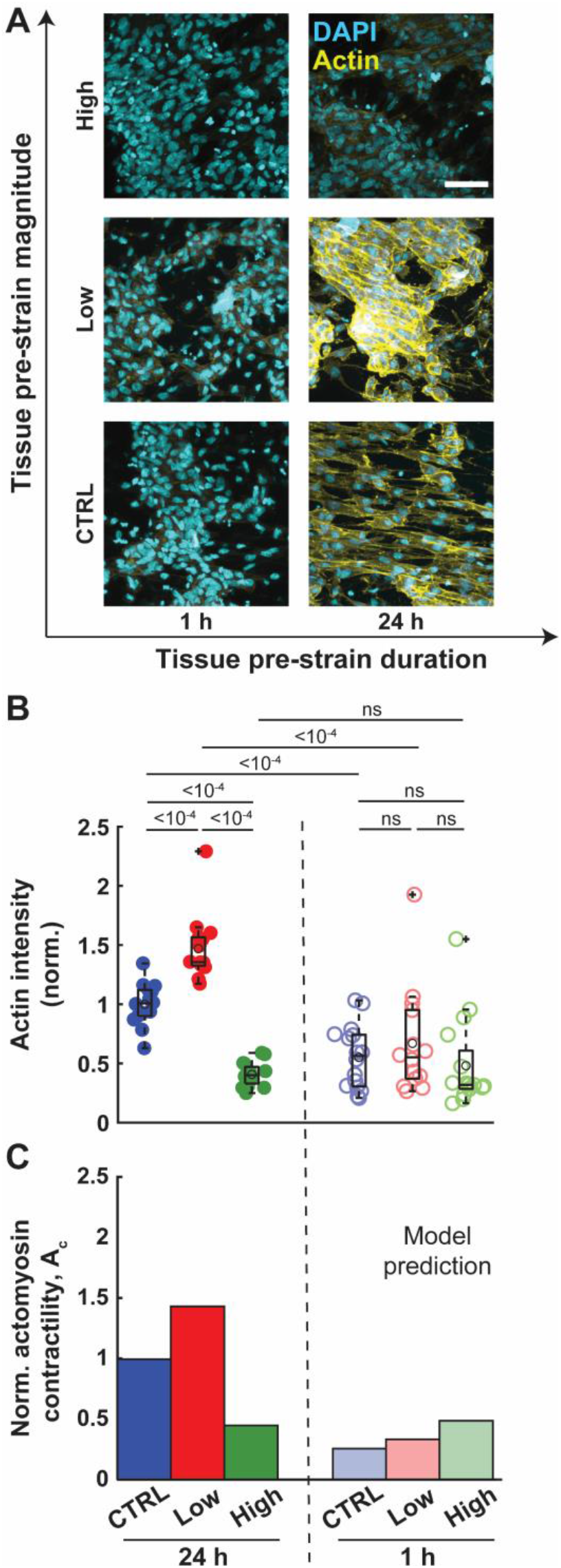
Stretching of skin tissues for a long duration can induce mechanical memory in cells and cause their long-term activation in a non-monotonic manner with tissue strain magnitude. (A-B) To determine the long-term level of cell activation, tissues were fixed 1 day after tissue unloading and the expression levels of F-actin in fibroblasts per cell were measured. F-actin is a well-known indicator of fibroblast activation level. Scale bar = 50 μm. *n* ≥ 14 for both 24h and 1h groups. (C) The cell activation level induced by the magnitude and duration of tension is predicted by the signaling network model.

### Long-term tissue contraction force depends on both cell activation level and matrix stiffness after tissue unloading

A seeming contradiction is that cells in tissues stretched for 24 h in control or low stretch conditions had higher sustained levels of activation than cells in tissues stretched for 1 h (Fig. 5), but tissue-level contractile forces for these tissues stretched for 24 h were very close to those for tissues stretched for 1 h (Fig. 1E). Our theoretical framework identified that the mechanism underlying this is again the transmission of force from cells by the ECM. In the tissues stretched for 24 hours, the ECM softens (Figs. 2E,F), reducing the tissue-level stress associated with cell contraction. This is shown in Fig. 1F where we used our theoretical framework to determine the tissue contractile stress (*σ*) 1 day after tissue unloading (and not during the tissue stretching as in Fig. 4C). Model predictions showed good agreement with our experiments (Fig. 1E), and revealed that elevated cell activation and cell contractility do not necessarily lead to elevated tissue contraction because ECM stiffness modulates the transmission of cellular forces. The ECM softening associated with long-duration stretching attenuates the transmission of cell contractile forces to the tissue level (Fig. 4G-I).

## Conclusions

Using an integrated experimental and theoretical approach, we described mechanisms by which cell-ECM feedback modulates both the long-term activation of cells and tissue contractility. This occurred at strain levels known to be important for wound healing and skin grafting. Unlike previous observations on stem cells, fibroblasts developed mechanical memory under static tissue strain, depending on the duration and magnitude of tissue strain. Central to this was a role of the ECM in the development of mechanical memory in fibroblasts and their long-term activation, modulated by time- and strain-dependent physical properties of the ECM.

## Materials and Methods

### Cell culture and 3D tissue construction

Human dermal fibroblasts (HDFs), derived from adult dermis, were purchased from Lonza (Basel, Switzerland, Catalog #: CC-2511). We used cells only at passages P4 to P8 in our experiments. Cells were first cultured at 37°C with 5% CO_2_ in Dulbecco’s Modified Eagle Medium (DMEM) containing 10% fetal bovine serum (FBS) and 1% penicillin-streptomycin (PS). Cells were next detached with 0.05% Trypsin/EDTA for 5 min, separated with Accutase (Sigma Aldrich, St. Louis, MO), and resuspended in fresh DMEM. The suspended cells in DMEM were mixed with rat tail-derived type I collagen in 0.1% acetic acid and 2X DMEM (Sigma). After neutralizing the pH with sodium hydroxide, 1 mL of the mixture containing 1.5×10^6^ cells/mL HDFs and 1.2 mg/mL collagen was added into Teflon-made cylindrical casting molds with a mandrel at the center with a diameter of 9.65 mm (Fig. 1A). The tissues were incubated at 37 °C with 5% CO_2_ for 2 days and formed compacted and thin contractile rings with cells aligned in the circumferential direction of the molds.

### Tissue force measurement experiments

We measured the contractile force in each tissue before ((*F*^0^)_pre_: Fig. 1C(i)) and after ((*F*^0^)_post_: Fig. 1C(iv)) tissue stretching. The tissue force measurements were performed using a custom-made tensile tester with sensitive isometric force transducers (Harvard Apparatus) and stepper motors (Haward Industry) plugged into an Arduino for applying displacement precisely (Fig. 1B). A MATLAB subscript was developed for collecting the output voltages reported by the transducers, which were calibrated to the force. To control the temperature, a water pump with a temperature control system was connected to the organ bath (Fig. 1B).

The tissue rings were removed from the casting molds after 2 days of compaction (Fig. 1A) and were hung onto the attachment bars, bathed with 25 mM HEPES buffered DMEM with 10% FBS and 1% PS at 37 °C (Fig. 1B). The attachment bars were then separated at a 13.3 mm distance under the control of the Arduino (Fig. 1C(i)). At this length, the periphery of the tissues on the stretcher device is equal to the periphery of the tissues in the mold on day 2. The tissue force was measured by lifting the lower attachment bars away from the samples to define the zero-force point and bring it back to the 13.3 mm distance. Fig. S2 shows a representative plot of tissue force measurement. The average of forces measured during the first 200 s of force measurement was reported as the tissue contractile force.

After measuring the initial tissue contractile force, the tissues were moved back to the incubator and were kept in sodium bicarbonate buffered DMEM at 37 °C with 5% CO_2_. We designed 3D-printed spacers with different lengths and placed them between the two ends of the tissues to generate 0% (tissue length = 13.3 mm), 5%, and 30% strains in the tissues (Fig. 1C(ii)).

The tissues were kept under the prescribed strain levels either for 1 hour or 24 hours (Fig. 1C(ii)). The spacers were then removed and the tissues were allowed to relax for 1 day (Fig. 1C(iii)). We then measured the contractile force in each tissue using the force measurement procedure described above (Fig. 1C(iv)).

### Stress relaxation test

After measuring the contractile force of pre-strained tissues (Fig. 1C(iv)), tissues were allowed to relax for 10 minutes and then were hung onto the attachment bars of the tensile tester. We then applied 5% strain to the tissues and kept them under the strain for 10 minutes (Fig. 2A(v)) while we continuously measured and recorded the force (Fig. 2B). The stiffness of the tissue was measured as ΔF/ΔX where ΔF is the difference between forces measured at t = 10 min and t = 0, respectively, and ΔX = (0.05)(13.3 mm) is tissue deformation (Fig. S3).

We then allowed the tissues to relax for 10 minutes and treated them with 2 μM cytochalasin-D for 1 hour to disrupt the cell-generated contractile forces (Fig. 2A(vi)). We then repeated the stress relaxation test (Fig. 2A(vii)) and measured the stiffness as described above (Fig. 2E).

### Matrix stress-strain relationship

After the stress relaxation test on cytochalasin-D treated tissues, we allowed the tissues to relax for 10 minutes on the tensile tester. Starting from the initial tissue length of 13.3 mm, we then stretched the tissues with a loading rate of 3.11 μm/s to determine the force-strain curves in Fig. 3D. Note that, as depicted in Fig. 2E, tissues pre-strained for 24 h showed ECM softening, while tissues pre-strained for 1h did not exhibit permanent changes in their long-term ECM stiffness. Therefore, for the determination of the matrix stress-strain relationship depicted in Fig. 3D, we only used tissues pre-strained for 1 hour.

### Immunofluorescence and cell imaging in 3D collagen gels

To determine the long-term level of cell activation, we fixed the tissues one day after tissue unloading and stained them for F-actin (Figs. 5A-B) and α-SMA (Fig. S4A-B). Tissues were fixed with 4% paraformaldehyde (Electron Microscopy Sciences, Catalog #: 15714) for 2 hours at 37°C, permeabilized with 0.5% triton X-100 solution (Sigma, X100) for 30 minutes and blocked with 10% goat serum (Thermos Fisher, Catalog #: 50062Z) for 1 hour at room temperature before incubating with anti-α-smooth-actin antibody (1:250 dilution, Abcam, Catalog #: ab124964) in 10% goat serum at 4°C overnight. Tissues were incubated with rhodamine-phalloidin (1:400 dilution, Thermos Fisher, Catalog #: R415) with Alex Fluor 488-conjugated secondary antibody (1:500 dilution, Abcam, Catalog #: ab150077) in 10% goat serum for 2 hours at room temperature. Each step was followed by PBS washing for a minimal 10 minutes at least 3 times. Samples were mounted to glass-bottom dishes with a prolonged gold antifade mounting solution with DAPI (Thermos Fisher, Catalog #: P36931). Z-stack images were obtained through a Zeiss LSM 880 laser confocal microscope with identical parameters in each experiment.

### Skin graft strain field

We used the Holzapfel-Gasser-Ogden model (SI 3) to determine the average strain magnitude in stretched skin grafts (Fig. S1). The model was first fitted to unmeshed skin stretching experiments from reference^48^ to determine the model parameters. We then used the calibrated model to simulate the stretching of a meshed skin graft with meshing ratios of 1.3, 3, and 5, commonly used in skin graft surgeries.^49^ The model parameters were shown in Table S3.

### Active chemo-mechanical tissue model and cell signaling network model

Details of the models were presented in SI 1 and SI 2, respectively. The code and a full description of model parameters are available on GitHub (https://github.com/Farid-Alisafaei/Stretch-Induced-Cellular-Memory.git).

## Notes

https://github.com/Farid-Alisafaei/Stretch-Induced-Cellular-Memory.git

### Competing Interest Statement

The authors have declared no competing interest.

### Summary of Updates

Adding new data

