## Supplementary Information for "Matrix Softening Controls Stretch-Induced Cellular Memory and Fibroblast Activation"

### Active chemo-mechanical tissue model and cell signaling network model

To understand mechanisms underlying the effects of pre-strain duration and magnitude on the long-term contractility of cells and tissues, we developed a computational framework that couples an active chemo-mechanical tissue model with a cell signaling network model.

### 1. Active chemo-mechanical tissue model

The active chemo-mechanical tissue model comprises three components; an active force-generating element and two passive elements (Fig S5). The active element represents fibroblast cells that generate active contractile forces and drive tissue deformation. The two passive elements correspond to the extracellular matrix (ECM) that underlies tissue architecture and biomechanical properties. One of these passive elements, which is connected in parallel with the active element, represents the family of ECM fibers that undergo compression as the cell contracts. Conversely, the other passive element, which is linked in series with the active element, reflects the family of ECM fibers that undergo tension as the cell contracts (Fig S5).

#### 1.1. Matrix in tension

The family of ECM fibers that is under tension, denoted as “matrix in tension” in Fig S5, is represented by a spring element and a viscous element. These two elements are considered in the model based on two key characteristics of the ECM observed in our experiments when subjected to tensile loading.

The first was that the mechanical properties of the ECM undergo long-term, time-dependent changes when subjected to sustained tensile strain. This observation was made during our stress-relaxation experiments, wherein tissues treated with the cell actomyosin inhibitor cytochalasin D were subjected to constant and sustained tensile strains. Our experimental results showed a gradual decrease in tension within the ECM over time, exemplified by Fig. 3C. This behavior illustrated the viscous nature of the ECM when subjected to tension. To capture this ECM behavior, our model incorporates a viscous element (Fig. S5). The stress within this element is related to the rate of ECM strain, which can be expressed as:

$$\sigma_\eta = \eta \dot{\varepsilon}_\mathrm{m} \tag{S1}$$

Here, $\eta$ represents the viscosity of the ECM, approximated to be 170 kPa/hr based on the stress relaxation tests conducted.

The second was that ECM stiffness changes with the magnitude of applied tensile strain as observed in our experiments in Fig. 3D. This was achieved through the stretching of tissues that had been treated with the cell actomyosin inhibitor, cytochalasin D. To minimize the effect of viscous behavior, we used a slow rate of stretching 3.11 µm/s, while simultaneously measuring the resulting force as a function of tensile strain (see Fig. 3D). By dividing the force by the cross-sectional area of the tissue (approximately 1 mm × 0.15 mm), we obtained the stress-strain curve for the ECM under tension. To incorporate this stress-strain relationship into the constitutive model of the spring element, we employed a smooth function (Eq. S2) which was fitted to the experimental stress-strain curve. The resulting fitted curve (shown as the dashed line in Fig. 4B) was subsequently used as the stress-strain relationship for the spring element in the model.

$$\sigma_\mathrm{s} = \begin{cases} B e^{\frac{\xi_1}{\eta_1}\tan^{-1}\left(\frac{\eta_1}{\xi_1}\right)} \sqrt{1+\left(\frac{\xi_1}{\eta_1}\right)^2}\, e^{-\xi_1(\varepsilon_2-\varepsilon_\mathrm{m})} \sin(\eta_1(\varepsilon_2-\varepsilon_\mathrm{m})), & \varepsilon_\mathrm{m} \le \varepsilon_\mathrm{t} \\ B e^{\frac{\xi_2}{\eta_2}\tan^{-1}\left(\frac{\eta_2}{\xi_2}\right)} \sqrt{1+\left(\frac{\xi_2}{\eta_2}\right)^2}\, e^{-\xi_2(\varepsilon_\mathrm{m})} \sin(\eta_2\, \varepsilon_\mathrm{m}), & \varepsilon_\mathrm{m} > \varepsilon_\mathrm{t} \end{cases} \tag{S2}$$

In Eq. (S2), $\varepsilon_\mathrm{m}$ is the strain in the ECM element in tension as shown in Fig. S5. All parameters in Eq. (S2), including $\eta_1 = \pi/\varepsilon_2$, $\xi_1 = \eta_1 \cot(\eta_1\varepsilon_1)$, and $\xi_2 = \eta_2 \cot[\eta_2(\varepsilon_2 - \varepsilon_1)]$, are expressed in terms of $\varepsilon_1 = 0.2$ and $\varepsilon_2 = 0.67$, which were determined by fitting Eq. (S2) to the stress-strain curve in Fig. 3D. Note that $\varepsilon_\mathrm{t} = \varepsilon_2 - \varepsilon_1$ is the point at which the stress starts to decrease. Also, $\eta_2 = \pi/\varepsilon_4$ where $\varepsilon_4 = 1.1$ is the strain at which rupture occurs. The maximum stress generated in the matrix, denoted as $B$ in Eq. (S2), was determined to be 40 kPa. This value was obtained by dividing the force measured from our uniaxial stretching experiments (Fig. 3D) by the corresponding cross-sectional area as described earlier.

The total stress experienced by the ECM element in tension, which is equal to the tissue stress $\sigma$ (Fig. S5), is determined by the summation of stresses in the spring and viscous elements.

$$\sigma = \sigma_\mathrm{s} + \sigma_\eta \tag{S3}$$

#### 1.2. Matrix in compression

The contraction of cells leads to tissue shrinkage, consequently causing a substantial portion of ECM fibers to undergo compression. To account for this group of fibers, we incorporated a spring element in parallel with the cell element in our model (Fig. S5). As a result, in the absence of external strain, this element undergoes compression due to the contraction of the cell element. To maintain simplicity, we represented the spring element as a linear and time-independent material. Subsequently, the stress in this element was determined as $E_\mathrm{c}\, \varepsilon_\mathrm{c}$ where $E_\mathrm{c}$ and $\varepsilon_\mathrm{c}$ are the stiffness and strain of the ECM element in compression, respectively. In our simulations, we used $E_\mathrm{c}$ = 9 kPa as measured in our previous work using atomic force microscopy (AFM) indentation experiments.[1]

#### 1.3. cell

The active contractile behavior of fibroblast cells is captured in the model by an active element that generates contractile stress, denoted as $\sigma_c$. This cell model is designed based on the observation that actomyosin contractility of fibroblasts, and consequently their contractile stress $\sigma_c$, increases in response to tension $\sigma$ present within the tissue, which can be generated by the inherent contractility of the cells themselves, as well as by externally applied tensile stress. To accomplish this, the total free energy of the cell, $W$, encompasses a combination of chemo-mechanical terms that account for the underlying processes involved.[2]

$$\mathrm{W}(\sigma_c, \varepsilon_c) = \underbrace{\sigma_c\, \varepsilon_c}_{\text{Cell work}} + \underbrace{\frac{\beta}{2}(\sigma_c - \sigma_c^0)^2 - \alpha \int_0^{\sigma_c} \sigma \, \mathrm{d}\sigma_c}_{\text{Chemical energy}} + \underbrace{\frac{1}{2} E_c\, {\varepsilon_c}^2 - \int_0^{\varepsilon_c} \sigma \, d\varepsilon_c}_{\text{Mechanical energy}} \tag{S4}$$

In Eq. (S4), the parameter $\alpha$ serves as a feedback parameter governing the increase of actomyosin contractility in response to tension. A higher value of $\alpha$ signifies a stronger feedback mechanism, promoting an increase in the contractile stress $\sigma_c$ in response to tension $\sigma$. Conversely, the parameter $\beta$ acts as a chemical stiffness parameter that attenuates the feedback mechanism. Consequently, a higher value of $\beta$ makes it more difficult for the cell to surpass its baseline contractility $\sigma_c^0$, thus weakening the amplification of $\sigma_c$ in response to tension. The values for the model parameters can be found in Table S1 estimated from our previous work.[1] Minimization of the total free energy with respect to the two unknown variables $\sigma_c$ , and $\varepsilon_c$ yields the following equations

$$\frac{\partial \mathrm{W}(\sigma_c, \varepsilon_c)}{\partial \varepsilon_c} = \sigma_c + E_c\, \varepsilon_c - \sigma = 0 \tag{S5}$$

$$\frac{\partial \mathrm{W}(\sigma_c, \varepsilon_c)}{\partial \sigma_c} = \varepsilon_c + \beta(\sigma_c - \sigma_c^0) - \alpha\sigma = 0 \tag{S6}$$

Eq. (S5) shows the interplay between cell stress $\sigma_c$, cell strain $\varepsilon_c$, and the stress generated in the tissue $\sigma$. In the absence of external strain, the stress $\sigma_c$ generated by the contraction of the cell element induces a compressive stress $E_c\, \varepsilon_c$ within the parallel ECM element, while simultaneously leading to a tensile stress $\sigma$ within the tissue.

$$\sigma_c = -E_c\, \varepsilon_c + \sigma \tag{S7}$$

By substituting the expression for $\sigma$ from Eq. (S6) into Eq. (S7), we can obtain the following equation that depicts the relationship between cell stress and cell contraction

$$\sigma_c = \frac{E_c\, \alpha - 1}{\beta - \alpha} \varepsilon_c + \frac{\beta}{\beta - \alpha} \sigma_c^0 \tag{S8}$$

where $\varepsilon_c$ is the cell strain. Note that in the absence of external strain, $\varepsilon_c \leq 0$ as the cell contracts. Also, it should be noted that cell strain is equal to the strain of the ECM element in compression, as these two elements are connected in parallel (Fig. S5).

As previously discussed, the design of the cell model is rooted in experimental observations indicating that fibroblast actomyosin contractility, and thus the cell-generated stress, intensifies in response to tension present within the tissue. This relationship becomes evident when substituting the expression for $\varepsilon_c$ from Eq. (S7) into Eq. (S8), resulting in the following equation

$$\sigma_\mathrm{c} = \frac{E_\mathrm{c}\,\alpha - 1}{E_\mathrm{c}\,\beta - 1}\sigma + \frac{E_\mathrm{c}\,\beta}{E_\mathrm{c}\,\beta - 1}\sigma_\mathrm{c}^0 \tag{S9}$$

which demonstrates the increase in cell stress ($\sigma_\mathrm{c}$) with tissue stress ($\sigma$). Also, Eqs. (S8) and (S9) show the range of cell stress in the absence of external strain; $E_\mathrm{c}\,\beta\sigma_\mathrm{c}^0/(E_\mathrm{c}\,\beta - 1) \leq \sigma_\mathrm{c} \leq \beta\sigma_\mathrm{c}^0/(\beta - \alpha)$. The minimum cell stress is obtained when the tissue is unconstrained and free to contract without any resistance ($\sigma = 0$). The maximum cell stress occurs when the tissue faces maximum resistance to contraction, achieved by immobilizing both ends of the cell ($\varepsilon_\mathrm{c} = 0$).

#### 1.4. Coupling of the cell and matrix models

The final equation needed to complete the set of equations can be derived from the following relationship (Fig. S5)

$$\varepsilon = \varepsilon_\mathrm{c} + \varepsilon_\mathrm{m} \tag{S10}$$

which states that the tissue strain $\varepsilon$ is the sum of the cell strain ($\varepsilon_\mathrm{c}$) and the strain in the ECM element connected in series ($\varepsilon_\mathrm{m}$). Note that $\varepsilon$ is known and varies from 0 to 0.3 in our simulations as measured in stretched skin grafts (Fig. S1) and used in our experiments (Fig. 1C). By incorporating this equation, the set of six unknown variables $\sigma_\eta$, $\sigma_\mathrm{s}$, $\sigma$, $\sigma_\mathrm{c}$, $\varepsilon_\mathrm{c}$, and $\varepsilon_\mathrm{m}$ can be determined by solving the complete set of equations comprising (S1)-(S3) and (S7)-(S9).

#### 1.5. Stability criterion

Note that $\beta$ and $\alpha$ cannot be equal, as their equality ($\beta = \alpha$) would result in the cell stress $\sigma_\mathrm{c}$ in Eq. (S8) and subsequently the overall stress $\sigma$ approaching infinity, which is not physically feasible. Additionally, the parameter $E_\mathrm{c}$ representing the elastic modulus of the ECM element in compression must always be positive, as negative values are not physically meaningful.

To ensure that $\sigma_\mathrm{c}$ is always positive, signifying actomyosin contractility pulling on the extracellular matrix rather than pushing, the coefficients $(E_\mathrm{c}\,\alpha - 1)/(\beta - \alpha)$ and $\beta/(\beta - \alpha)$ in Eq. (S8) should be positive. These constraints provide the following stability criterion:

$$\beta > \alpha > \frac{1}{E_\mathrm{c}}$$

#### 1.6. ECM stiffness

In a tissue stretched and held isometrically at strain $\varepsilon$, the ECM is stretched in tension by both cell-generated contractile stresses ($\sigma_\mathrm{c}$) and the external stresses applied to maintain the tissue strain of $\varepsilon$ (Fig. 4A). However, due to the viscous nature of the ECM, the tensile strain in the matrix, $\varepsilon_\mathrm{m}$, increases with tissue stretching duration (Fig. 4A). As a result, for long stretching durations (e.g., 24 h), the strain in the ECM can locally exceed the strain associated with peak stress in Fig. 3D. This effect could be seen in our simulations: after 1 h of isometric tissue stretching, the ECM strain stayed sufficiently low that the slope of the stress-strain curve ($E_\mathrm{tangent}$) was independent of tissue strain magnitude $\varepsilon$, but after 24 h, the ECM strain reached or passed the peak so that $E_\mathrm{tangent}$ dropped and varied with $\varepsilon$ (Fig. 4B).

To validate the model predictions against our experiments in Fig. 2E, we calculated the ECM stiffness ($k$) as shown in Fig. 2F. To this end, we converted the ECM modulus $E$, which has the unit of Pa=N/m, to $k$ which has the unit of N/m using $k = E\,A\,/\,L$ where $A$ = (1 mm)(0.1 mm) = 0.1 mm$^2$ is the tissue cross-section area and $L$ = 13.3 mm is the tissue length.

For tissues pre-strained for 1 h (short duration), the ECM remained elastic and did not show any permanent softening. Thus, the ECM modulus $E$ for these tissues can be calculated as the slope of the stress-strain curve ($E_{\mathrm{tangent}}$). For tissues pre-strained for 24 h (long duration), the ECM showed inelastic behavior and was softer compared to the ECM of tissues pre-strained for 1 h (Fig. 2E). Therefore, the ECM modulus *E* for 24 h pre-strained tissues was calculated as $E = (1 - D)\, E^o$ where *D* is the inelastic (damage) parameter and $E^o$ is the ECM elastic (undamaged) modulus (i.e., $E_{\mathrm{tangent}}$ in tissues pre-strained for 1 h).

The inelastic (damage) parameter *D* increases linearly with the matrix strain, $\varepsilon_{\mathrm{m}}$, until it reaches its maximum value *D* = 1 at $\varepsilon_{\mathrm{m}} = \varepsilon_{\mathrm{r}}$ which is the matrix strain at which rupture occurs. This can be captured by the following equation

$$D = D_1 + \frac{\varepsilon_{\mathrm{m}} - \varepsilon_1}{\varepsilon_2 - \varepsilon_1}(D_2 - D_1)$$

where $\varepsilon_2 = \varepsilon_{\mathrm{r}}$ is the matrix strain at which rupture occurs and $\varepsilon_1 = \varepsilon_{\mathrm{m}}^{\mathrm{CTRL,24h}}$ is the matrix strain in the tissue pre-strained for 24 h with 0% strain. Also,

$$D_2 = 1 \quad , \quad D_1 = 1 - \frac{E_{\mathrm{m}}^{\mathrm{CTRL,24h}}}{E^o}$$

where $E_{\mathrm{m}}^{\mathrm{CTRL,24h}}$ is the slope of the stress-strain curve ($E_{\mathrm{tangent}}$) in the tissue pre-strained for 24 h with 0% strain.

### 2. Signaling network model

The active chemo-mechanical tissue model serves to determine the stress experienced by cells during tissue pre-straining. The signaling network model serves to determine how the stress field that cells experience during pre-straining affects the long-term activation of cells which persists after the tissues have been unloaded.

The signaling network model was developed based on a known cascade of biological processes (i) to (viii) described in the main text and represented in Fig. 4D. Each of these biological processes (i) to (viii) was translated into a corresponding mathematical representation shown in Eqs. (S11)-(S14).

The input to the model was the cell-generated contractile stress $\sigma_{\mathrm{c}}$ (stress that cells experienced during tissue stretching) determined from the active chemo-mechanical tissue model (Fig. S6). The model outputs were the time variation of four normalized variables: (1) the normalized actomyosin activation level, $A_{\mathrm{c}}$; (2) the normalized level of translocation of cell activation factors to the nucleus, $T_{\mathrm{c}}$; (3) the normalized cell-level expression of genes *not* associated with the myofibroblastic phenotype, $g$; and (4) the normalized cell-level expression of genes associated with the myofibroblastic phenotype, $G$ (Fig. 4D). These were determined by solving the following set of equations.

First, tensile stress promoted cell actomyosin contractility through pathways such as Rho-Rock or activation of mechanosensitive ion channels (Fig. 4D i).[2–5] This is represented in the model by incorporating the terms denoted as (i) in Eq. (S11), where the cell stress $\sigma_{\mathrm{c}}$ increases the cell actomyosin activation level $A_{\mathrm{c}}$, using a Hill function format.

$$\frac{dA_{\mathrm{c}}}{dt} = k_1 \underbrace{\frac{(\sigma_{\mathrm{c}}/K_1)^{n_1} + (g/K_2)^{n_2}}{1 + (\sigma_{\mathrm{c}}/K_1)^{n_1} + (g/K_2)^{n_2}}}_{\text{(i), (ii)}} + k_2 \underbrace{\frac{(\sigma_{\mathrm{c}}/K_3)^{n_3} + (G/K_4)^{n_4}}{1 + (\sigma_{\mathrm{c}}/K_3)^{n_3} + (G/K_4)^{n_4}}}_{\text{(i), (iii)}} - d_1 A_{\mathrm{c}} \qquad \text{(S11)}$$

Elevated levels of cell contractility trigger the translocation of transcriptional and epigenetic factors, such as MKL and YAP/TAZ, into the nucleus (Fig. 4D iv).[6] This is represented by incorporating the term denoted as (iv) in Eq, (S12), where the cell actomyosin activation level $A_{\mathrm{c}}$ increases the translocation of cell activation factors to the nucleus $T_{\mathrm{c}}$.

$$\frac{dT_{\mathrm{c}}}{dt} = \underbrace{k_3\, A_{\mathrm{c}}}_{\text{(iv)}} - d_2\, T_{\mathrm{c}} \tag{S12}$$

The enhanced nuclear accumulation of cell activation factors subsequently stimulates the transcription of genes associated with actomyosin contractility and cell activation (Fig. 4D vi, vii).[7,8] To model this process, the terms denoted as (vi) and (vii) are incorporated into Eqs. (S13) and (S14), respectively.

$$\frac{dg}{dt} = k_4 \underbrace{\frac{(A_{\mathrm{c}}/K_5)^{n_5}}{1 + (A_{\mathrm{c}}/K_5)^{n_5} + (T_{\mathrm{c}}/K_6)^{n_6}}}_{\text{(v), (vi)}} - d_3\, g \tag{S13}$$

$$\frac{dG}{dt} = k_5 \underbrace{\frac{(T_{\mathrm{c}}/K_7)^{n_7}}{1 + (A_{\mathrm{c}}/K_8)^{n_8} + (T_{\mathrm{c}}/K_7)^{n_7}}}_{\text{(vii), (viii)}} - d_4\, G \tag{S14}$$

In these equations, $T_{\mathrm{c}}$ downregulates the transcription of genes *not* associated with cell activation ($g$), while simultaneously upregulating the transcription of genes associated with cell activation ($G$), such as genes involved in actin-cytoskeleton.[7] Subsequently, expression of these genes increases cell actomyosin contractility which is captured in the model through a feedback loop (Fig. 4D ii, iii), represented by the terms (ii) and (iii) in Eq. (S11).

Finally, note that while increased levels of cell contractility stimulate the translocation of activation factors into the nucleus, subsequently promoting the transcription of genes related to actomyosin contractility and cell activation, excessive cell contraction can result in cell detachment from the extracellular matrix, as shown in our previous work.[9] When cells detach from the extracellular matrix, the tension that was previously generated at the cell-matrix interface diminishes entirely, resulting in a substantial reduction in their actomyosin contractility. Consequently, this decrease in contractility can trigger a transition in the cells, leading to a transcriptionally less active state (Fig. 4D v, viii).[8] This transition is captured within the model through the terms (v) and (viii) in Eqs. (S13) and (S14), respectively. Specifically, an elevation in the cell actomyosin activation level ($A_{\mathrm{c}}$) leads to the upregulation of genes *not* associated with cell activation ($g$), while simultaneously downregulating the transcription of genes directly involved in cell activation ($G$).

### 3. Skin graft strain field

We used the Holzapfel-Gasser-Ogden (HGO) model in the Finite Element Software, Abaqus, to determine the strain field in stretched skin grafts. To determine the model parameters, we first fitted the stress-strain curves, determined from uniaxial skin stretching simulations along and parallel to the Langer line, to unmeshed skin stretching experiments from reference.[10] Fig. S1D showed the uniaxial stretching simulations of unmeshed skin grafts harvested parallel and perpendicular to the Langer line (Fig. S1B). The model parameters are shown in Table S3.

Subsequently, the calibrated model was used to simulate the stretching of meshed skin grafts. In these simulations, a skin graft with an initial area of $A = ab$ was meshed using meshing ratios of 1.3, 3, and 5, which are commonly used in skin graft surgeries (Fig. S7).[11] The skin meshing ratios were calculated using the following standard equation[12,13]

$$R_{\mathrm{n}} = 1 + \left[\frac{1}{d} \times \frac{1}{L+g} \times \left(\frac{L}{2}\right)^2\right] \tag{S15}$$

where $L$ is the length of the cut, $g$ is the distance between the cuts along the cut direction, and $d$ is the distance between the cuts perpendicular to the cut direction (Fig. S7). The meshed skin grafts were stretched perpendicular to the direction of cuts to expand horizontally and cover a wound with a predefined area of 1.25*A*. This was done in our simulations by moving the right edge of the graft $1.85a$ horizontally (Fig. S1E). As expected, the simulations showed a nonuniform distribution of strain, with higher strain values observed around the vertices of the cuts (Fig. S1E). The average value of the maximum principal strain was plotted in Fig. S1E for each meshing ratio.

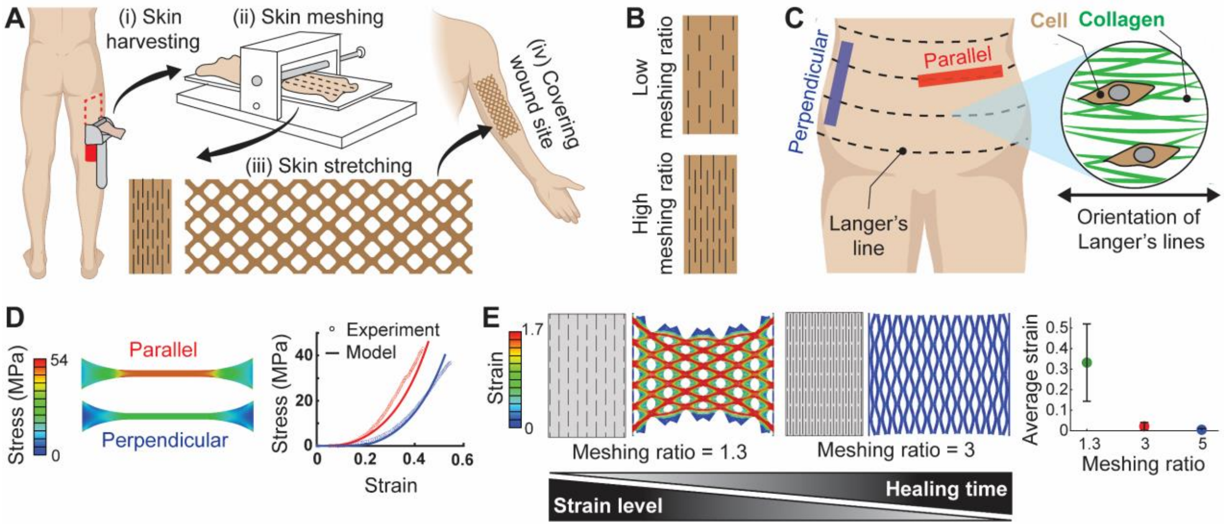


Figure S1. The strain level that skin tissues experience during skin graft stretching can be tailored by an order of magnitude by tuning skin graft meshing ratio. (A) Skin grafting exemplifies a medical procedure in which skin tissues undergo long-term external stretching. To protect from pathogens, large chronic wounds are often treated with split-thickness skin grafts in which epidermis and part of the dermis are harvested from a donor site (i), then cut into a meshwork of slits (ii) and stretched (iii) over a larger wound site (iv). (B) Schematics of a skin graft meshed with different meshing ratios. The choice of meshing ratio impacts the strain field that cells experience in the stretched skin graft. (C) Skin is an anisotropic material with orientation-dependent mechanical properties where fibroblast cells and collagen fibers are mainly aligned along topological lines called Langer's lines. (D) To determine how the strain field in skin grafts changes with meshing ratio, we first calibrated a hyperelastic model against uniaxial stretching experiments of unmeshed skin grafts. (E) We then used the calibrated model to determine how the strain field changes with meshing ratio. The simulations showed the strain fields generated in skin tissues depend strongly on the design of slits with the choice of meshing ratio affecting the average strains in graft tissue by an order of magnitude. (Strains and stresses in D and E represent the maximum principal strain and stress, respectively. The details and the parameters of the hyperelastic model used in our simulations in D and E can be found in Materials and Methods, SI 3, and Table S3. Skin stretching experiments in D were taken from reference[10]. In panel E, the solid circles and the bars represent the mean value and standard deviation, respectively.)

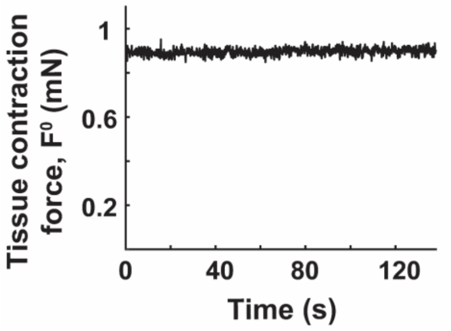


Figure S2. A representative illustration of the measurement process for tissue contraction force, as used in Fig. 1. All force measurements were conducted at the tissue's original length of 13.3 mm. The tissue contraction force was monitored over a period of 200 seconds, and the average force value recorded during this 200-second interval was selected as the tissue contraction force.

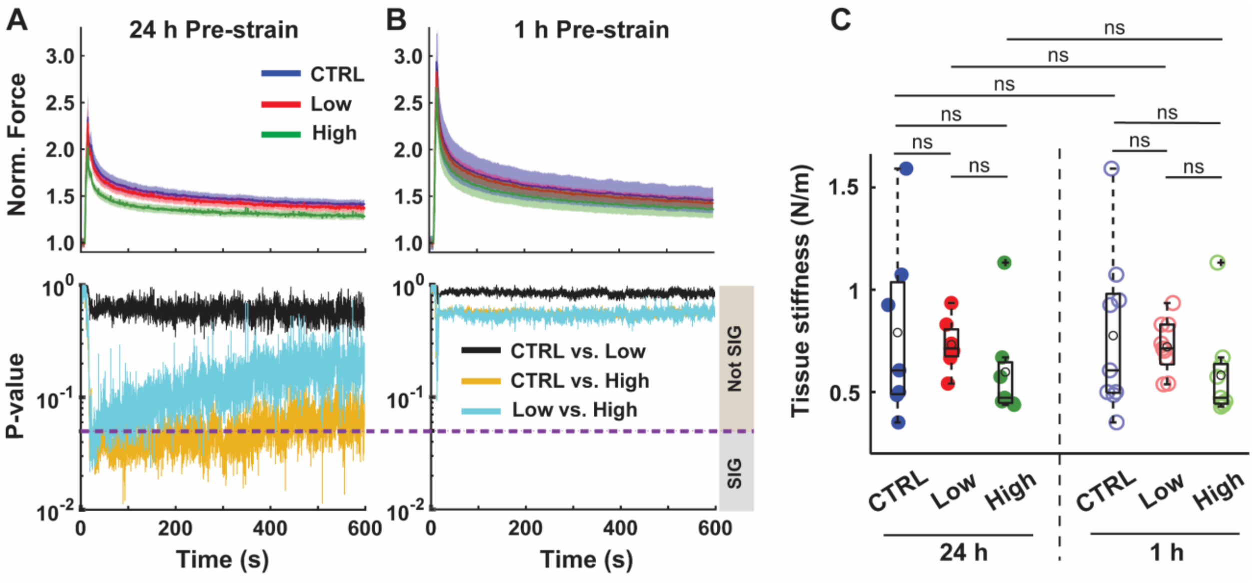


Figure S3. Effects of pre-strain magnitude and duration on the long-term responses of tissues were quantified via stress relaxation tests after the completion of tests described in Fig. 1. (A-B) Stress relaxation tests were conducted on the tissues pre-strained for 24 h and 1 h, in which tissues were stretched by an amount $\Delta X$=0.66 mm and held at that length while the force was recorded. The force rose to a peak during the stretching, and then relaxed to an asymptotic value over time. The difference $\Delta F$ between the pre-test isometric force (force measured at 0 s) and the force measured after 600 s of relaxation was recorded, and (C) the long-term tissue stiffness was calculated as $\Delta F/\Delta X$. In A and B, the solid lines and the shaded areas represent the mean and standard error, respectively. *n* = 7 and 9 for the 24h and 1h groups, respectively.

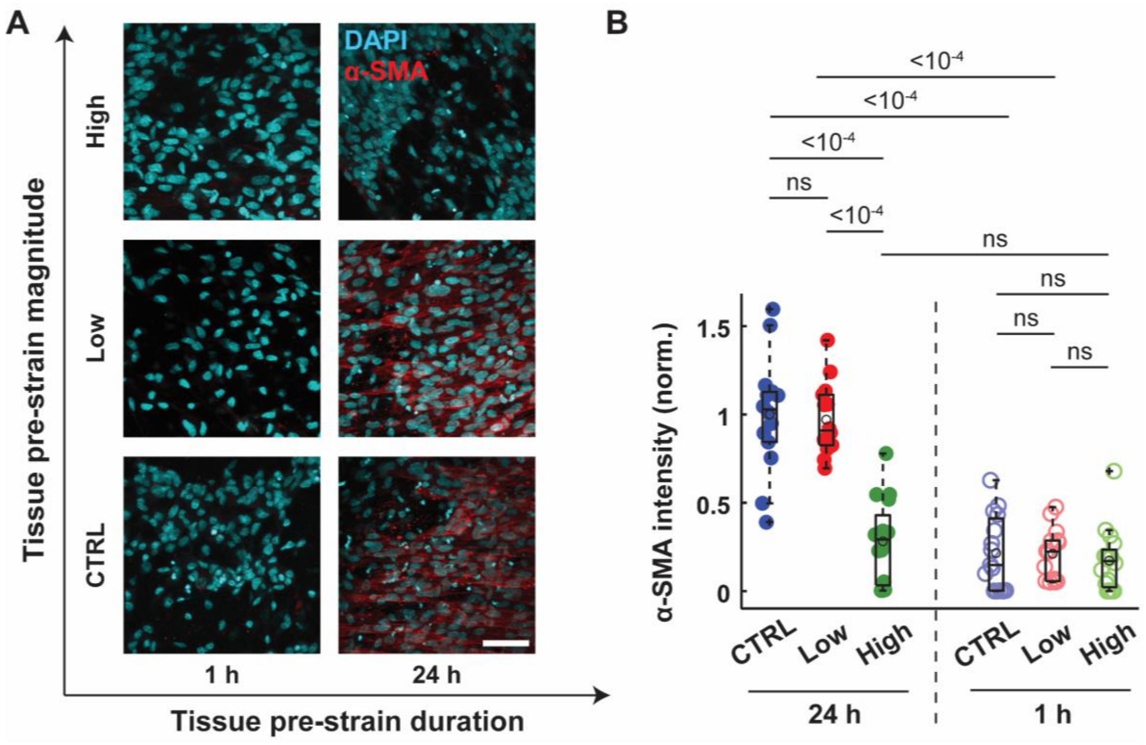


Figure S4. To determine the long-term effect of pre-straining magnitude and duration on the level of cell activation, tissues were fixed 1 day after tissue unloading and the expression levels of α-smooth muscle actin (α-SMA) per cell were measured. The results showed that cell activation occurred only in tissues pre-strained at 0% (CTRL) and 5% (Low) for 24 h. Scale bar = 50 µm. *n* ≥ 14 for both 24h and 1h groups.

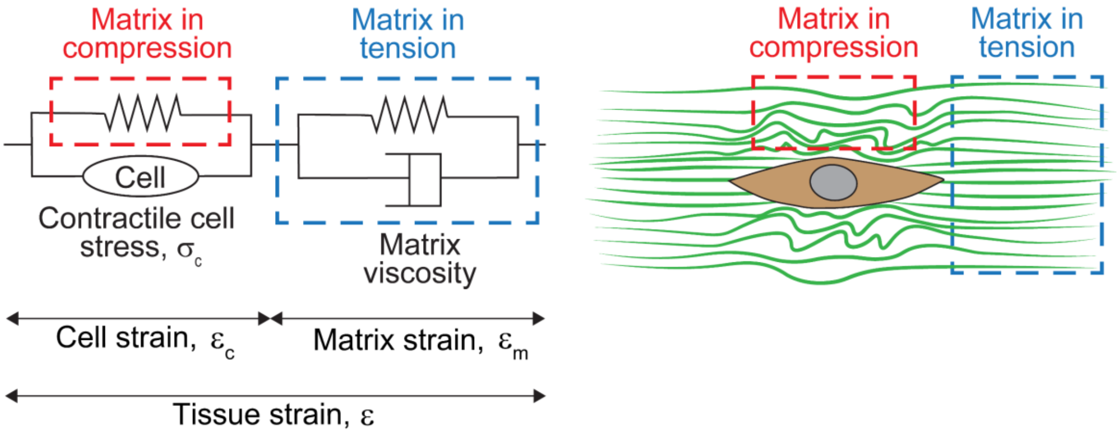


Figure S5. Graphical representation of the active chemo-mechanical tissue model comprising an active force-generating element (cell) linked to two passive elements (matrix). In the absence of external strain ($\varepsilon = 0$), the matrix element connected to the cell element in parallel represents regions of the extracellular matrix experiencing compression due to cell contraction, while the series matrix element represents regions of the ECM under tension.

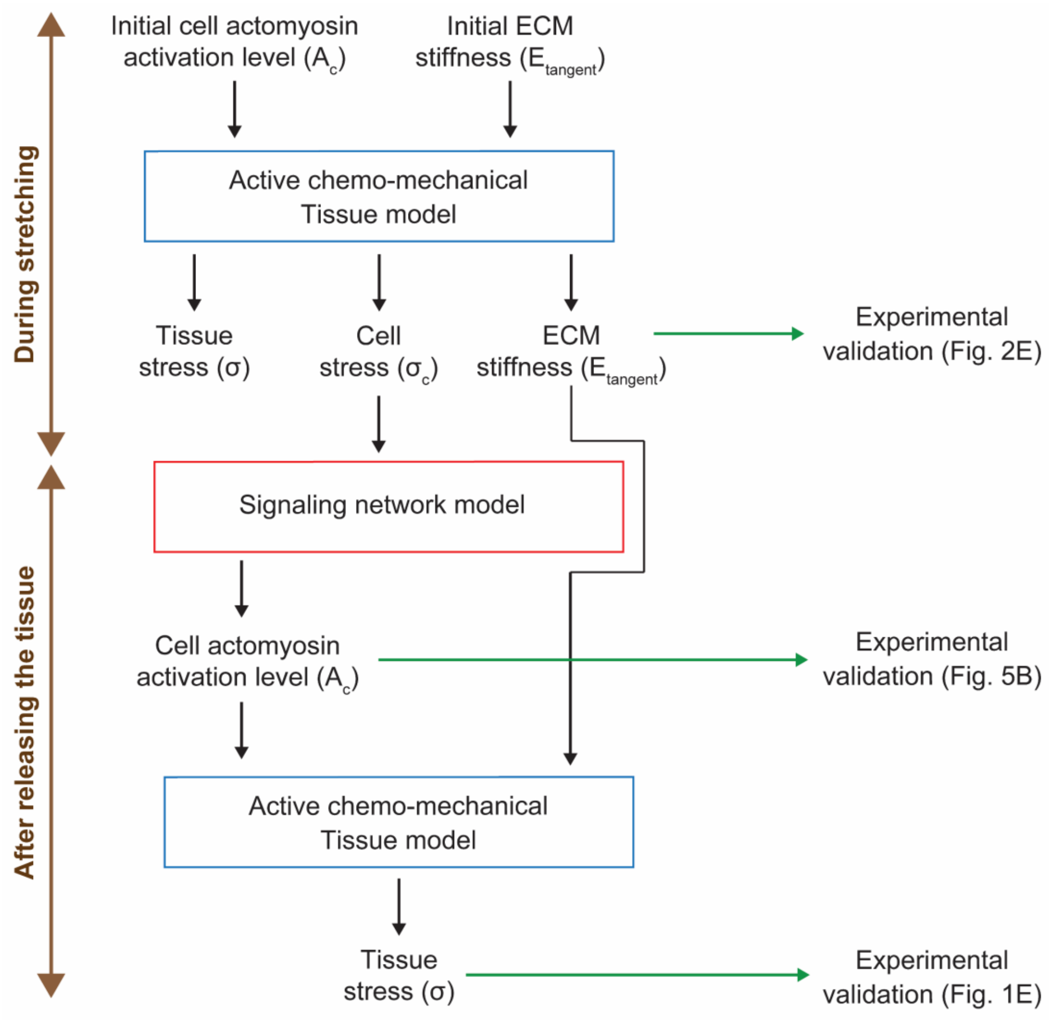


Figure S6. Flowchart illustrating the integration of the active chemo-mechanical tissue model with the cell signaling network model, along with corresponding model predictions and experimental validations.

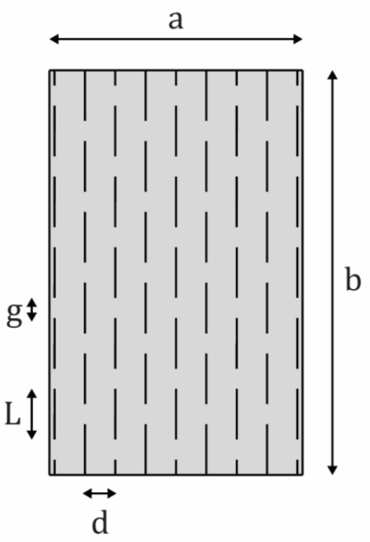


Figure S7. Schematic representation of the three skin grafting parameters $L$, $g$, and $d$ used in Eq. S15.

Table S1. Model parameters of the active chemo-mechanical tissue model.

| Parameter | Description | Value | Unit |
|---|---|---|---|
| $\sigma_c^0$ | Baseline contractility | 6.2 | kPa |
| $\alpha$ | feedback parameter | 0.45 | 1/kPa |
| $\beta$ | chemical stiffness parameter | 0.5 | 1/kPa |

Table S2. Model parameters of the cell signaling network model.

| Parameter | Value | Parameter | Value | Parameter | Value |
|---|---|---|---|---|---|
| $K_1$ | 1.1 | $n_1$ | 55.4 | $k_1$ | 0.3 |
| $K_2$ | 110 | $n_2$ | 4.4 | $k_2$ | 90 |
| $K_3$ | 100000 | $n_3$ | 3.6 | $k_3$ | 4 |
| $K_4$ | 1.5 | $n_4$ | 38 | $k_4$ | 10 |
| $K_5$ | 1 | $n_5$ | 5 | $k_5$ | 1 |
| $K_6$ | 0.5 | $n_6$ | 8 | $d_1$ | 0.15 |
| $K_7$ | 160 | $n_7$ | 2.5 | $d_i$ (i=2,3) | 1 |
| $K_8$ | 45 | $n_8$ | 5.5 | $d_4$ | 0.8 |

Table S3. Model parameters used for the skin graft stretching simulations in Fig. S1.

| $C_{10}$ | $k_1$ | $k_2$ | D | $\kappa$ |
|---|---|---|---|---|
| 0.1007 MPa | 24.53 MPa | 0.1327 | 0 | 0.1535 |
